# Cultivation and characterization of human midbrain organoids in sensor integrated microfluidic chips

**DOI:** 10.1101/869701

**Authors:** Sarah Spitz, Cristian Zanetti, Silvia Bolognin, Mudiwa Nathasia Muwanigwa, Lisa Smits, Emanuel Berger, Christian Jordan, Michael Harasek, Jens C. Schwamborn, Peter Ertl

## Abstract

With its ability to emulate microarchitectures and functional characteristics of native organs *in vitro*, induced pluripotent stem cell (iPSC) technology has enabled the generation of a plethora of organotypic constructs, including that of the human midbrain. However, reproducibly engineering and differentiating such human midbrain organoids (hMOs) under a biomimetic environment favorable for brain development still remains challenging. This study sets out to address this problem by combining the potential of iPSC technology with the advantages of microfluidics, namely its precise control over fluid flow combined with sensor integration. Here, we present a novel sensor-integrated platform for the long-term cultivation and non-invasive monitoring of hMOs under an interstitial flow regime. Our results show that dynamic cultivation of iPSC-derived hMOs maintains high cellular viabilities and dopaminergic neuron differentiation over prolonged cultivation periods of up to 50 days.

## 2. INTRODUCTION

With the emergence of iPSC technology, it has now become possible to emulate complex cytoarchitectures and key characteristics of native tissues *in vitro*, including that of the human brain.^1^ In recent years so-called “organoids” have demonstrated their potential as powerful tools for modeling key aspects in organ development, homeostasis as well as pathology.^2–4^ Efforts have been made to establish iPSC-derived models of the human midbrain, that recapitulate organotypic characteristics including spatial organization, neuronal activity as well as disease-associated phenotypes.^5,6^

However, the inability of conventional culture techniques, to precisely control and reproduce cellular microenvironments has led to issues with organoid variability, among others, a key limitation of three-dimensional (3D) technology.^7^ In addition, common strategies fail to take biophysical cues, such as e.g. interstitial flow, into account, fundamental aspects in organ development and maturation.^7^ Interstitial fluid flow not only plays an important role in the delivery of nutrients and removal of metabolic waste, but it implicates non-synaptic cell-cell communication, ionic homeostasis, cell migration as well as immune function. In addition, interstitial fluid flow is involved in drug delivery, distribution and clearance and thus of considerable importance when considering organoid models in the context of drug screening applications.^8^ Overall, the lack of biophysical and other environmental cues restricts the development of physiologic organoids in a more complete and reproducible manner, specifically important when emulating structures of high intrinsic complexity such as those of the human brain.^7^

Microfluidics with its inherent ability to provide precise spatial and temporal control over cellular microenvironments coupled with the capability of non-invasive monitoring (e.g. embedded microsensors), low reagent consumption, and increased throughput provides an ideal tool for tackling said aspects.^8^ Organ-on-a-chip approaches were already shown to beneficially impact cerebral organoid differentiation and alter pathological responses in an *in vitro* Alzheimer model under an interstitial flow regime, underlining its potential in neurobiological applications.^13–15^ Up to now, however, microfluidic based organoid models focusing on neurobiological aspects are limited and still fail to include non-invasive monitoring strategies, an important tool for addressing the issue of organoid variability. Here, we investigated a novel strategy for differentiating iPSC-derived hMOs in sensor-integrated microfluidic platforms under dynamic conditions by directing interstitial fluid flow via hydrostatic pressure through the growing organoid. Computational fluid dynamic (CFD) simulations were employed to characterize flow characteristics and subsequently ensure low shear stresses as well as interstitial flow regimes within the microfluidic device. Non-invasive oxygen monitoring was established to asses organoid viability and growth, while maturation was confirmed by immunohistochemical analysis.

## 3. METHODS

### Chip Fabrication

Microstructures were fabricated by soft lithography from a CNC milled mold using polydimethylsiloxane (PDMS) (Sylgard® 184 Silicone Elastomer Kit, Down Corning). After polymerization at 80 °C, molded PDMS was bonded to glass substrates using air plasma (Harrick Plasma, High Power, 2 min). As previously described, microfluidic devices equipped with oxygen sensors were generated by the deposition of microparticles into drilled cavities within glass substrates by the use of a pipette. After drying for 2 h at room temperature, the microparticles were immobilized to the glass substrate and the fluidic structures were sealed, employing air plasma.^9^ Prior to use microfluidic devices were sterilized employing 70 % ethanol as well as UV plasma.

### Finite Volume CFD Simulation

A multipurpose finite volume CFD code (Ansys Fluent 6.3.26, www.ansys.com / OpenFoam www.openfoam.org) was used for solving the flow problem. The geometry consisting of the hydrogel cavity, the two feed channels as well as the two collection units was split into 136.000 hexahedral control volumes. The grid pillars at the gel inflow and outflow boundary have been fully resolved. For adequate numerical accuracy, second or higher order discretization schemes have been selected for all flow variables (Navier-Stokes equation – momentum conservation, Continuity equation – mass conservation) and for the species equations. All wall boundaries were treated as ideally smooth; no slip boundary conditions (zero flow velocity at the wall) were selected for all surfaces. The outlet was set to pressure outlet at a standard pressure of p = 1 atm (101325 Pa). The hydrogel region was approximated as homogeneous and isotropic porous zone (Darcy-Forchheimer equation) with a constant porosity of ε = 0.99 and a viscous resistance of R = 1.33·10^13^ 1/m² having been assumed for all directions.^16^

Isothermal flow was assumed, no temperature or energy field was solved. For simplicity, Newtonian fluid behavior was applied for the simulation using a constant dynamic viscosity and constant density (incompressible) for all of the mixture components. As the concentrations of the dissolved species in the fluid are low, the properties of the solvent, water, have been used for the simulation (ρ = 993 kg/m³, η = 0.001003 Pa·s at 37 °C). The diffusion coefficients for the tracer components have been estimated according to literature values (glucose: 0.18 kDa – 4·10^−10^ m²/s, oxygen: 32 Da – 2·10^−9^ m²/s, water: 18 Da – 2·10^−9^ m²/s) assuming a dilute solution.^11^ To investigate the cross mixing of the two inlet channel fluids, different water species have been used for both inlets. Simulations were carried out on the cluster server cae.zserv.tuwien.ac.at (operated by the IT department of TU Wien, www.zid.tuwien.ac.at). As the major flow resistances are inside the hydrogel and in the flow channels, but not in the feed and collection cavities, a simplification was used: To reduce the computational effort, steady state simulations for different selected feed cavity filling levels have been carried out. The simulated filling level was translated into a corresponding relative pressure difference between feed inlet zone and pressure outlet.

### Fluorescein Assay

To validate data obtained from the CFD simulation a fluorescein assay was performed. For this purpose, fluorescein (Sigma Aldrich) was added to the medium reservoirs and its propagation within the microfluidic device was monitored using a Live Cell microscope (Pecon). Images were analyzed using ImageJ (N = 4).

### hMO Generation and On-Chip Cultivation

iPSCs from one healthy individual were used in this study. The maintenance of iPSCs was performed as previously described.^5^ From the iPSC line human ventralized neural epithelial stem cells (hvNESC) were generated, which were subsequently used to generate midbrain organoids.^5^ For on-chip hMO cultivation, organoids suspended in Matrigel^®^ (Corning^®^) were transferred into the microfluidic chip on day 0 of maturation phase and cultivated for up to 50 days. Dynamic cultivation was achieved by filling the feed medium reservoirs up to a 3 mm feeding level, while medium at the collector side was kept at 0.6 mm height. Medium was exchanged every 48 h.

### On-Chip Oxygen Monitoring

On-chip oxygen monitoring was carried out at a sampling frequency of 1 Hz using a FireStingO_2_ optical oxygen meter (Pyroscience) connected to optical fibers (length 1 m, outer diameter 2.2 mm, fiber ^diameter 1 mm). Integrated sensors were calibrated using a CO_2_/O_2_ oxygen controller (CO2-O2-^ Controller 2000, Pecon GmbH) equipped with integrated zirconium oxide oxygen sensors. Oxygen measurements were performed once a week (n=4). For this purpose, chips were sealed with PCR foil and transferred into an external incubation chamber setup. Samples were measured for 10 minutes to guarantee proper equilibration. Oxygen demand was subsequently calculated according to the following ^formula: hMO oxygen demand (∆P_O2_) = P_O2_ blank − P_O2_ hMO.^

### Viability Assay and Morphological Characterization of hMOs

To monitor the viability of hMOs a viability assay employing calcein-AM and ethidium homodimer-1 (Invitrogen, L3224) was performed every week (n=3). Images were analyzed using the ImageJ plugin Color Pixel Counter. Cellular viabilities were determined by dividing the number of green pixels by the total number of red and green pixels, while exposure time and focus plane were kept constant for each analysis. To obtain information on both hMO growth as well as neurite outgrowth, organoid diameter (highest diameter of each organoid) and neurite lengths were assessed manually using ImageJ.

### Immunofluorescence

hMOs were fixed with 4 % PFA and washed 3x with phosphate-buffered saline (PBS) for 15 min. They were embedded in 3-4 % low-melting point agarose in PBS. The solid agarose block was sectioned with a vibratome (Leica VT1000s) into 50 µm sections. The sections were blocked on a shaker with 0.5 % Triton X-100, 0.1 % sodium azide, 0.1 % sodium citrate, 2 % bovine serum albumin and 5 % normal goat serum in PBS for 90 min at room temperature. Primary antibodies were diluted in the same solution but with only 0.1 % Triton X-100 and were applied for 48 h at 4 °C. The following first antibodies were used: anti-rabbit Tuj1 (Optim AB Eurogentec, 1:600); anti-rabbit TH (SantaCruz, 1:1000).

After incubation with the primary antibodies, sections were washed 3x with PBS and incubated with the secondary antibodies in 0.05 % Tween-20 in PBS for 2h at RT and washed with 0.05 % Tween-20 in PBS and Milli-Q water before they were mounted in Fluoromount-G mounting medium (Southern Biotech). The organoids were imaged using a confocal microscope (Zeiss).

### Statistical Analysis

Data analysis was performed using GraphPad Prism software. A paired *t*-test was performed to determine significant differences between the individual data sets.

## 2. RESULTS

### Hydrostatic pressure driven flow-based microfluidic platform

To allow for the physiologic cultivation of hMOs on chip under an interstitial flow regime, a hydrostatic pressure driven flow-based design was selected (see Figure 1). The PDMS-based microfluidic chip contains three individual chambers, interconnected by two micropillar arrays. While the two outer chambers form the medium channels, the middle chamber is designed to accommodate the differentiating hMO in a three-dimensional matrix such as Matrigel. By adjusting the filling volumes on both sites of the microfluidic chip various hydrostatic pressures can be generated and subsequently direct medium flow of different velocities not only through the hydrogel matrix within the central channel, but also through the embedded organoid. Therefore, nutrient supply is not limited to diffusion, but nutrients are actively transported to the embedded organoid, while simultaneously keeping shear forces at a minimum.

**Figure 1:**
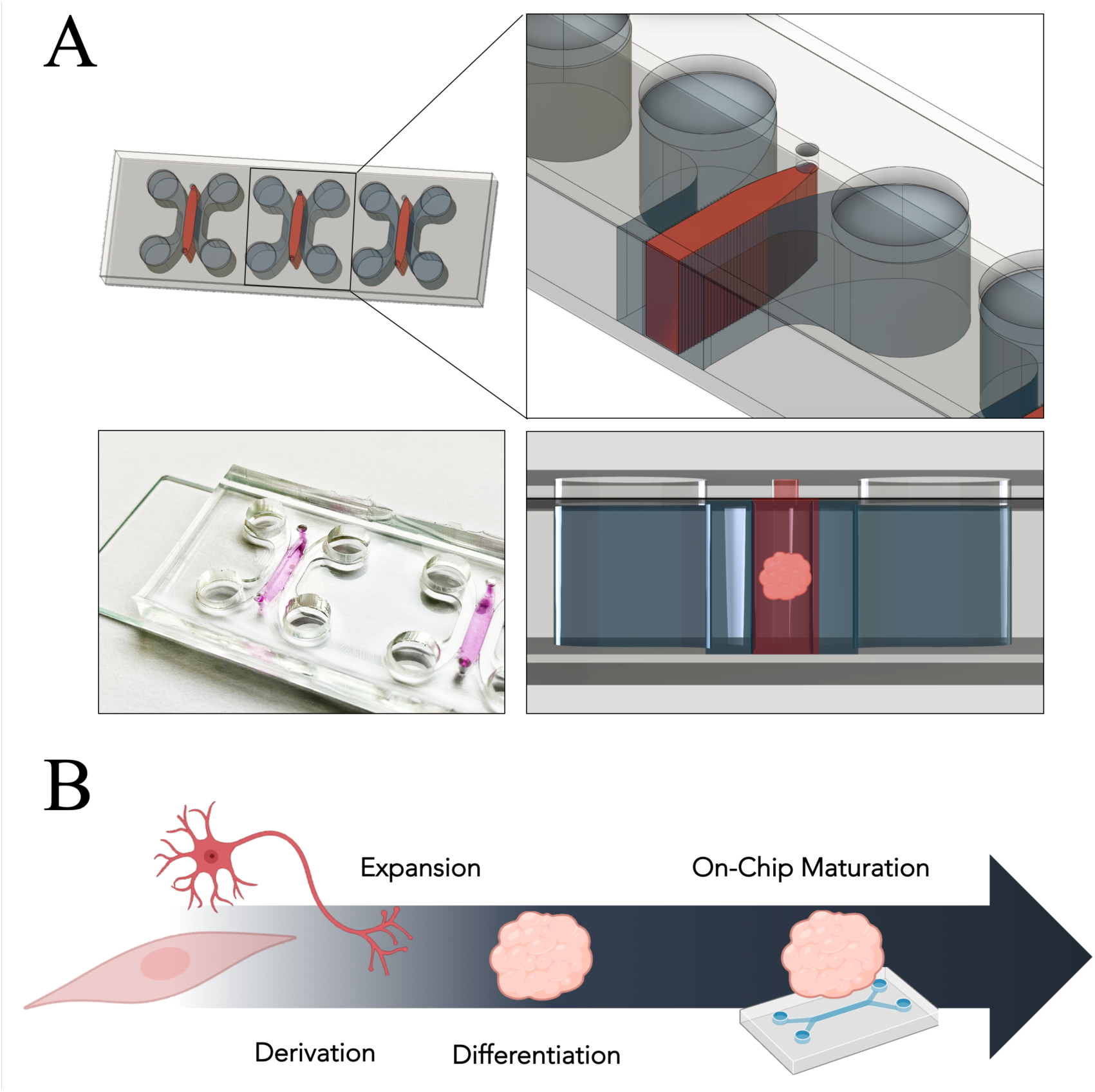
Hydrostatic pressure driven flow design comprising three channels interconnected by two micropillar arrays (A). Workflow of hMO cultivation. (B)

### CFD simulation of microfluidic platform reveals interstitial fluid flow

To determine optimal culture conditions and to assess overall flow characteristics within the microfluidic device CFD simulations were performed. The simulations were specifically set out to address platform specific correlations of overall volume flow rates and average velocity magnitudes within the central hydrogel chamber as a function of the relative inlet pressure.

For this purpose, the hydrostatic pressure driven flow-based design was split in hexahedral control volumes, with an example grid depicted in Fig. 2A, indicating the inflow and outflow boundary zones in blue and red, respectively. The flow within the microfluidic device was shown to behave strictly laminar, consequently resulting in a linear relationship between inlet pressure and volume flow rate. Initial assessment further revealed that the pressure drop within the microfluidic device can be mainly attributed to the hydrogel chamber, with negligible pressure loss resulting from the geometry of the device itself (see Figure 2B). As a result thereof, a proportionality function between flow rate and pressure drop was established, allowing for the calculation of flow velocity as a function of matrix permeability. Due to its beneficial properties to support self-organization of brain organoids, the CFD model and subsequent experiments were built upon the hydrogel matrix Matrigel, with a literature matched resistance value of R = 1.33·10^13^ 1/m² having been assumed for all directions.^16^ The simulated volume flow within the proposed hydrogel matrix of the microfluidic device was shown to behave highly uniformly throughout the most part of the central chamber. Parallelly aligned streamlines retrieved from the simulation at an exemplary level difference of 1 mm (equivalent to approx. 19 Pa pressure difference) confirm the aforementioned uniform velocity profile with an average velocity of 0.7 µm/s within the central part of the hydrogel chamber (see Figure 2C). While the comb structure of the micropillar array creates velocity gradients at the border of the hydrogel, the velocity within the hydrogel chamber, embedding the organoid, remains constant (see Figure 2D). Higher velocities close to the border zone also alter the velocity profiles and thus may induce shear stress on the embedded organoid. Since the occurrence, however, is restricted to both the upper and lower part of the hydrogel chamber flow inflicted shear stress on the organoid situated within the middle of the chamber can be ruled out (see Figure 2E). Overall, due to the uniform distribution of flow velocities within the central region of the hydrogel chamber and thus absence of velocity gradients, shear stresses within the central part of the chamber are kept at a minimum and thus generate optimal culture conditions for hMOs. Based on the established CFD model an initial reservoir pressure difference of 2.4 mm was selected to drive interstitial fluid flow through the hydrogel and thus provide optimal culture conditions for the organoid. By applying said hydrostatic pressure, hMOs were kept under dynamic culture conditions with flow velocities ranging from 1.6 µm/s (the upper limit of interstitial flow) down to 0.1 µm/s (the lower limit of interstitial flow) during the course of cultivation. Finally, to validate CFD simulation data, a fluorescein assay was employed. Fluorescein data were shown to be in good agreement with the calculated fluid regime with an initial measured average velocity of 1.67 ± 0.34 µm/s compared to an CFD estimated initial velocity of 1.6 µm/s.

**Figure 2:**
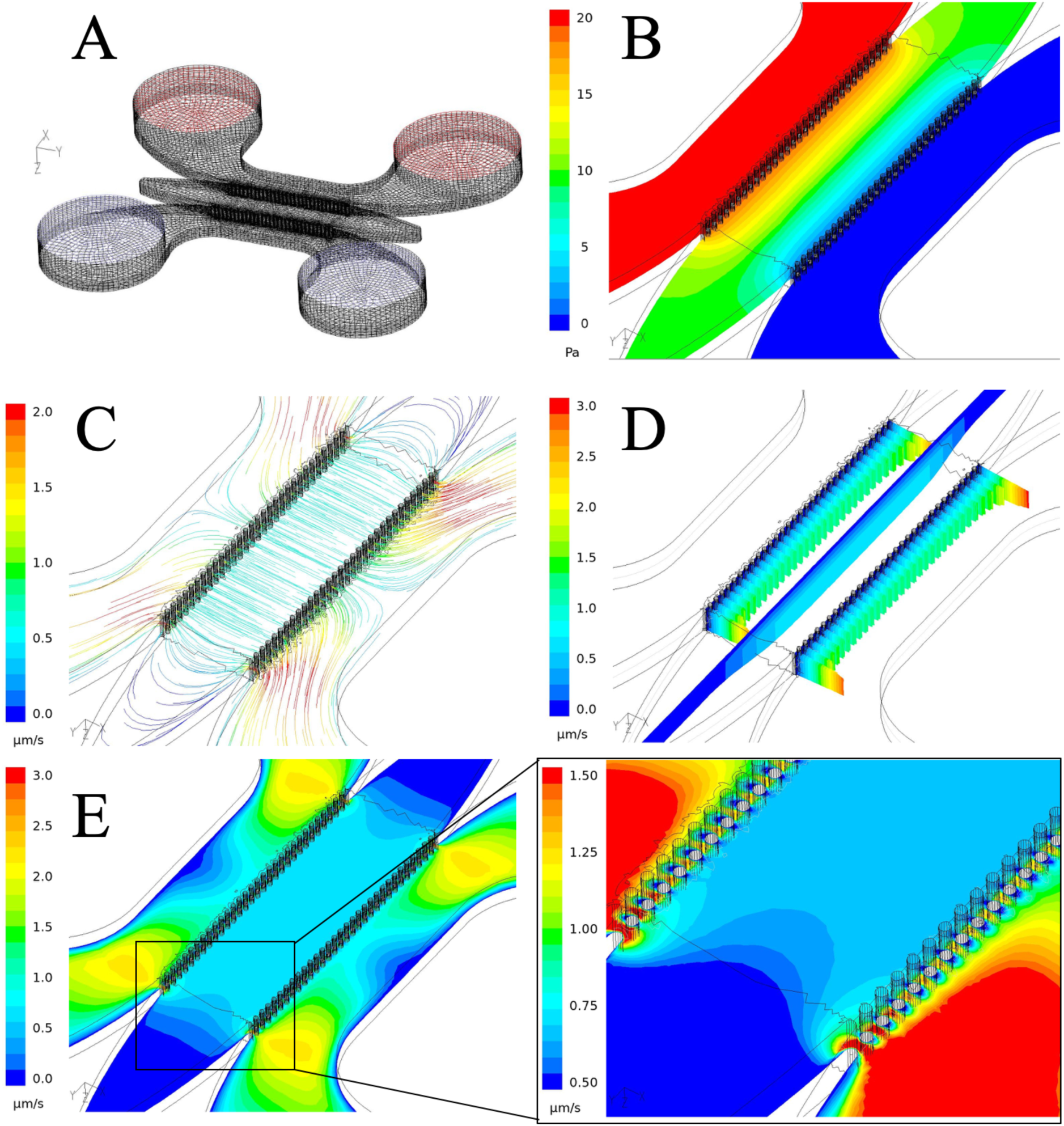
CFD simulation of the hydrostatic pressure driven flow design. Computational mesh of microfluidic device (A). Pressure gradient established within the central hydrogel chamber of the device (B). Streamlines with uniform flow velocities established at a pressure difference equivalent to 1mm level difference (C). Velocity profile in x-projection at the inlet, outlet and within the hydrogel chamber (D). Contour plot of velocity magnitude in symmetry plane of the microfluidic platform (E).

### Midbrain organoids differentiate in microfluidic chips

As an initial assessment of hMO behavior on chip neurite outgrowth, a crucial parameter for the formation of mature neural networks, was determined.^13^ After four days of cultivation under an interstitial flow regime hMOs displayed extensive neurite outgrowth, as seen in Figure 3A. While organoids cultivated for 24 h on-chip displayed an average neurite length of 293 ± 78 µm, hMOs cultivated for 96 h already displayed processes extending throughout large sections of the hydrogel chamber with an average neurite length of 1,024.4 ± 193 µm. Consequently, neurites extending from the embedded organoid display an average growth rate of 274.5 ± 26 µm/day, in good agreement with recently published microfluidic neural circuit models.^18^ Since midbrain organoid maturation is a time-consuming process with cultivation times ranging up to 100 days, it is of the utmost importance to guarantee prolonged cultivation without impairment of cellular viabilities. To ensure that hMOs remain viable within the proposed microfluidic device, organoids were analyzed using calcein AM/ ethidium homodimer-1 staining (see Figure 3B). Overall, no impairment in organoid viability was detected over the entire cultivation period of 6 weeks with an average viability of 93.8 ± 3.4 % (see Figure 3C). Furthermore, a significant increase in hMO diameter was detected after 42 days on chip (p < 0.001). Whereas hMOs displayed an average diameter of 737.4 ± 19.2 µm after one week of cultivation, the diameter increased up to 1,082.6 ± 66.2 µm after 6 weeks of culture within the microfluidic platform, indicative of organoid growth (see Figure 3C). In addition to an endpoint-based cell viability assay luminescent based oxygen sensor spots were integrated into the microfluidic device to ensure proper oxygenation as well as to non-invasively monitor hMO viability and growth (see Figure 3C). While the partial oxygen pressure measured within the blank chips remained constant over the entire cultivation period of 6 weeks with an average of 190.7 ± 16.7 hPa, thus ensuring sensor stability, a three-fold increase in oxygen demand could be detected in chips carrying hMOs. A small increase in oxygen demand was measured during the first three initial weeks of cultivation ranging from 51.6 ± 20.8 hPa up to 69.3 ± 23.8 hPa, followed by a significant increase by week 4 (p < 0.05) with an average oxygen demand of 92.5 ± 17.3 hPa which further increased up to 144.4 ± 17.1 hPa by week 6. Overall, the data indicate that hMOs cultivated within microfluidic devices remain viable for prolonged periods of time, while steadily increasing in size under dynamic culture conditions. These results corroborate well with aforementioned endpoint-based viability data. Furthermore, observed increase in oxygen demand may be linked to a metabolic switch from anaerobic glycolysis (exploited by hvNESC) to oxidative phosphorylation (predominant in differentiated neural cells), characteristic for organoid differentiation.^19^ Consequently, in a final set of experiments, maturation into midbrain specific organoids was assessed. For this purpose, hMOs dynamically cultivated for 30 days on chip were analyzed using immunohistochemical staining for the neuronal marker TUJ1 as well as for the dopaminergic neuron marker tyrosine hydroxylase (TH). Dynamically cultivated hMOs not only displayed robust differentiation into TUJ1-positive neurons, as shown in the histological section in Figure 3D (right panel), but in addition stained positive for TH (left panel).

**Figure 3:**
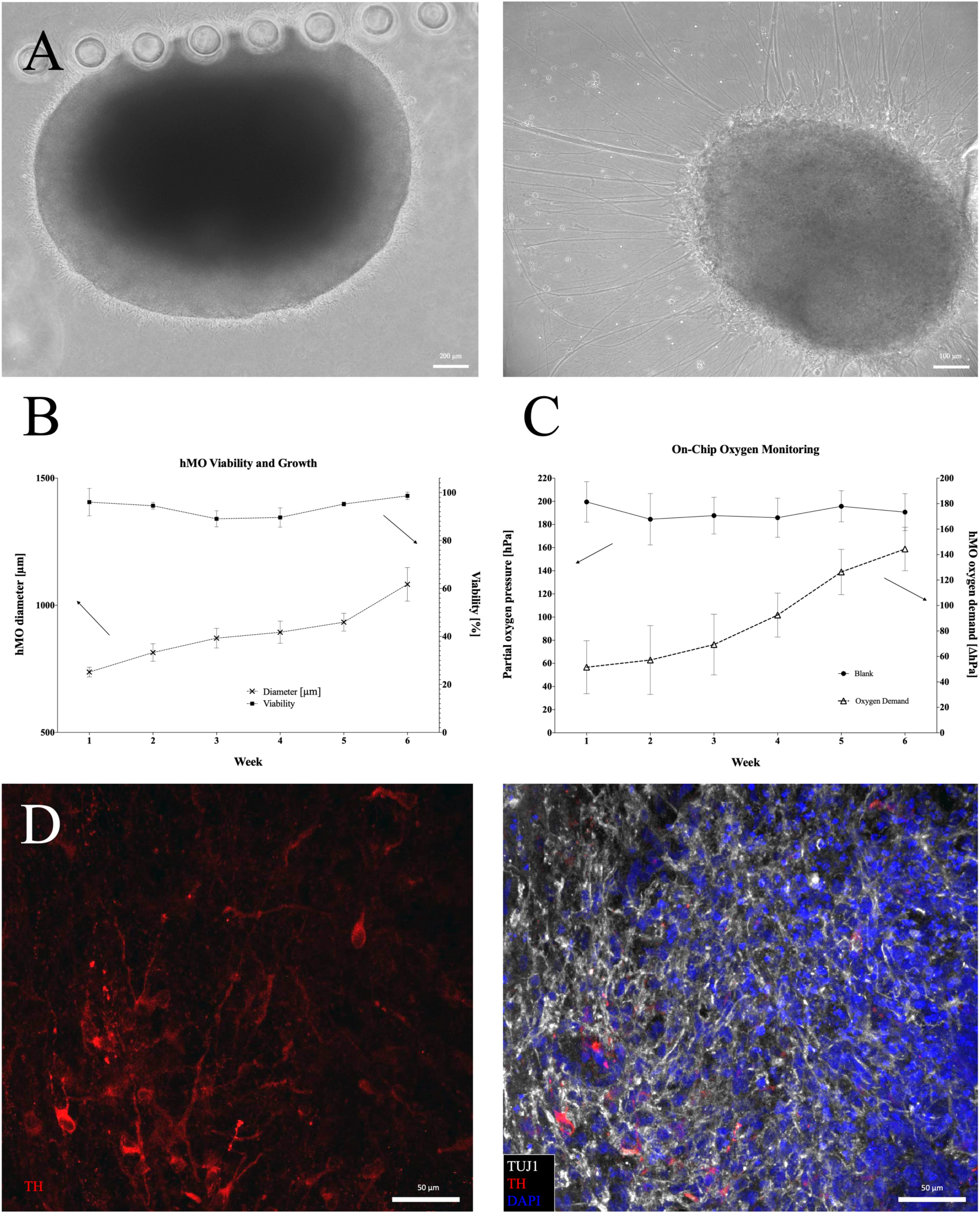
Brightfield images of hMOs after 24 h on chip and after 72 h revealing extensive neurite outgrowth (A). Time resolved graph showing hMO viability and organoid growth over a period of 6 weeks (n=3) (B). Time-resolved oxygen consumption of organoids on chip (n=4) (C). Immunohistochemical staining of hMO after 30 days on chip revealing TH (red) positive dopaminergic neurons as well as the neuronal marker TUJ1 (white) and DAPI stained nuclei (D).

## DISCUSSION

In this study, we developed a sensor-integrated microfluidic platform, which addresses not only the 3D cytoarchitecture of the human midbrain but in addition expands the model by the application of interstitial flow, a crucial parameter for nutrient transport, tissue maintenance as well as pathobiology. Overall, we showed that hMOs could be cultivated for prolonged cultivation periods of up to 50 days, without the impairment of cellular viability, essential for long term studies required in the field of 3D technology. These observations could be demonstrated, not only by the use of invasive endpoint assays but in addition were underlined by the integration of luminescent based oxygen sensors.

We have previously shown, that microfluidic culture can be beneficial for neuronal differentiation while simultaneously provide an economically efficient route to personalized drug discovery for Parkinson’s disease.^20^ However, the microfluidic cultivation of individual neuroepithelial stem cells in a 3D matrix, has resulted in poor cellular viabilities associated with high stress exposure during cell loading, limiting its use within the tested set-up.^6^ While our millifluidic approach has shown ameliorated differentiation behavior of hMOs combined with enhanced oxygen supply and a reduction in the necrotic core, the throughput remained limited.^19^ By combining hMOs with microfluidic technology we have not only addressed aforementioned limitations such as low viabilities and throughput but further improved aspects such as sensor integration, enabling non-invasive monitoring. Overall, our model integrates well with published midbrain-organoid literature: similar to published conventional culture set-ups, hMOs cultivated for 30 days within the presented microfluidic chip not only stained positive for the neuronal cell marker TUJ1 but also for the dopaminergic neuron marker TH.^5,6,21,22^ In summary, this study has provided a novel *in vitro* model that allows for prolonged organoid cultivation under an interstitial flow regime, favorable neurite outgrowth as well as non-invasive monitoring. The presented model is capable of emulating midbrain associated phenotypes in physiologic microenvironments that meet the need for relevant *in vitro* models and thus may provide a powerful tool in the context of midbrain pathologies such as Parkinson’s disease in the near future.

## 3. AUTHOR CONTRIBUTIONS

S.S. conceived, designed, and conducted experiments and analyzed data; S.B. and E.B. analyzed data and supervised the experiments; L.S. generated the hvNESC and hMOs; MNM performed the immunofluorescence experiments; C.J. and M.H. conducted CFD simulation. C.Z. conducted experiments. P.E. and J.S. supervised the project; S.S., P.E. and J.S. wrote the paper.

## 4. ACKNOWLEDGEMENTS

We thank Mrs. Thea van Wuellen and Mrs. Kyriaki Barmpa for excellent technical assistance. The JCS lab is supported by the Fonds National de la Recherché (FNR), Proof-of-Concept program PoC15/11180855 & PoC16/11559169) and M-era.Net project NanoPD (INTER/MERA/17/11760144 This is an EU Joint Programme-Neurodegenerative Disease Research (JPND) project (INTER/JPND/14/02; INTER/JPND/15/11092422). LMS was supported by a fellowship from the FNR (AFR, Aides à la Formation-Recherche). MNM is supported by the National Research Fund through the PARK-QC DTU FNR PRIDE17/12244779/PARK-QC.

